# A Maize Practical Haplotype Graph Leverages Diverse NAM Assemblies

**DOI:** 10.1101/2020.08.31.268425

**Authors:** Jose A. Valdes Franco, Joseph L. Gage, Peter J. Bradbury, Lynn C. Johnson, Zachary R. Miller, Edward S. Buckler, M. Cinta Romay

## Abstract

As a result of millions of years of transposon activity, multiple rounds of ancient polyploidization, and large populations that preserve diversity, maize has an extremely structurally diverse genome, evidenced by high-quality genome assemblies that capture substantial levels of both tropical and temperate diversity. We generated a pangenome representation (the Practical Haplotype Graph, PHG) of these assemblies in a database, representing the pangenome haplotype diversity and providing an initial estimate of structural diversity. We leveraged the pangenome to accurately impute haplotypes and genotypes of taxa using various kinds of sequence data, ranging from WGS to extremely-low coverage GBS. We imputed the genotypes of the recombinant inbred lines of the NAM population with over 99% mean accuracy, while unrelated germplasm attained a mean imputation accuracy of 92 or 95% when using GBS or WGS data, respectively. Most of the imputation errors occur in haplotypes within European or tropical germplasm, which have yet to be represented in the maize PHG database. Also, the PHG stores the imputation data in a 30,000-fold more space-efficient manner than a standard genotype file, which is a key improvement when dealing with large scale data.

## Introduction

The functional diversity of maize (*Zea mays* ssp. *mays* L.) makes it one of the most important crops, enabling its adaptation across the world (Hake & Ross-Ibarra, 2015). Maize is also genomically diverse, having gone through two rounds of whole-genome duplication, one ~70M years ago when grasses diverged (Paterson et al., 2004), and a second one ~11M years ago, an allotetraploidization event that was followed by diploidization (Gaut & Doebley, 1997). In addition to whole-genome duplication events, tremendous activity by transposable elements has further contributed to its genome diversity (Oliver et al., 2013). Previous studies have found considerable variability in the presence and absence of transcribed genes due to transposon activity (Fu & Dooner, 2002)(Lai et al., 2005). More recent analyses have leveraged whole-genome assemblies, allowing a detailed view of the extent of the intraspecific changes between maize varieties. These have identified signals of gene reordering, copy number, and other structural variations (Sun et al., 2018), with some cases accounting for 1.6Gb (equivalent to ~50% of the genome of B73) of variable transposable element sequences (Anderson et al., 2019).

Because maize is both a powerful system for genetics and evolutionary research, along with being a major crop worldwide, accounting for over 38% of the world’s cereal production (*FAO*, 2018), it has been genotypically characterized through different approaches that suited each community's needs and the technology available at the time. However, there is a tremendous need to leverage knowledge across these disciplines to facilitate breeding and understand the molecular and evolutionary basis of diversity. This paper leverages the recently assembled NAM founders and various types of sequencing data from thousands of maize lines to call high-density SNP genotypes at a unified set of sites.

The maize Nested Association Mapping (NAM) population was created as a resource for capturing a large proportion of broad maize diversity in a single population. It represents a highly studied set of ~5,000 recombinant inbred lines, generated from the single seed descent of an F1 crossing between 25 diverse maize inbreds and into a common parent, B73 (McMullen et al., 2009), allowing for the dissection of the genetic components underlying the control of maize phenotypes. The NAM population mapping resource (Buckler et al., 2009) has been utilized to identify quantitative trait loci for various complex traits (Gage et al., 2020). A recent NSF funded project (*NAM Genomes Project*, 2020) has produced extremely high-quality chromosome level assemblies of the NAM founders by combining long-read sequencing and optical mapping technologies. This is the first time the maize community has had a large set of equal quality assemblies. Here we aim to leverage these assemblies through a pangenome graph to impute the NAM population’s genotypes and other diverse germplasm.

Genotyping technologies vary in cost, accuracy, and number of sites and samples that can be genotyped in a single experiment (Romay, 2018). Specifically, the public maize community has used three major platforms: 1) Genotyping by Sequencing (GBS) has been used on tens of thousands of samples (Gouesnard et al., 2017; Rodgers-Melnick et al., 2015; Romay et al., 2013; Romero Navarro et al., 2017; Wu et al., 2016), but is challenged by short-read mapping issues and single-reference biases; 2) whole-genome sequencing (WGS) has been used in thousands of samples (Bukowski et al., 2018; Wang et al., 2020), with high variability in coverage and a similar reference mapping bias; and 3) SNP arrays, used to genotype thousands of samples over 55K and 600K sites (Unterseer et al., 2014; Xu et al., 2017), which by design have a predefined set of variant positions which are targeted to be genotyped. Additionally, new amplicon approaches (e.g., rhAmpSeq (Zou et al., 2020)) continue to be developed to increase the number of samples genotyped in a single experiment. All these distinct approaches represent a hurdle that needs to be addressed whenever there’s an interest in analyzing across genotyping experiments. Whole-genome imputation with a pangenome allows each of these technologies’ strengths and weaknesses to be complemented.

Imputation is the process of predicting genotypes that cannot be directly determined in a sample undergoing genotyping. In human studies, imputation approaches tend to leverage large reference panels (Browning et al., 2018; Das et al., 2016), composed of thousands to tens of thousands of samples with haplotypes identified through extremely dense SNP sets (Bycroft et al., 2018; McCarthy et al., 2016; Telenti et al., 2016). In the case of samples not represented by a reference panel, or when expecting some degree of relationship between the individuals in the sample, other imputation approaches attempt to identify identity-by-descent (IBD) segments from individuals that happen to have genotypes with higher marker density than the rest in the sample. The presence and identification of IBD segments allow un-genotyped SNPs’ imputation in lower density individuals by identifying their underlying haplotype. However, the human genome is an order of magnitude less diverse than plants like maize, and genotyping platforms for plants generally produce much less dense genotype marker sets. BEAGLE (Browning & Browning, 2013) is a leading tool in humans for within-sample imputation; it is also commonly used in crops, as it performs reasonably well in diverse and heterozygous populations with stable marker sets or high coverage (Chan et al., 2016; Pook et al., 2019). Other imputation approaches aim to leverage the peculiarities of populations within breeding programs; examples are FILLIN (Swarts et al., 2014), which utilizes breeding bottlenecks to capture libraries of haplotypes, and AlphaImpute (Hickey et al., 2012), which leverages the complex pedigrees of the samples under study to impute them.

To better capture the diversity of the plant genomes, some approaches represent this diversity as a collection of haplotypes in a graph, such as VG (Garrison et al., 2018). However, at present, VG cannot deal with the level of diversity in species such as maize. Another haplotype graph approach that can address this is the Practical Haplotype Graph (PHG) (Bradbury et al. in prep). In sorghum, a plant species more diverse than humans, the PHG was used to leverage haplotypes derived from parental samples, sequenced at high coverage, to impute progeny sequenced at very-low coverage (Jensen et al., 2020). Imputation in maize is challenging because of the high levels of divergence and repetitiveness result in poor read mapping. Also, its structural variation makes any single reference genome a poor model for the entire species. However, maize has an extensive collection of inbred varieties, where phasing of alleles becomes unnecessary. Through domestication, maize has gone through various selection bottlenecks, generating a modest subset of highly diverse haplotypes, as most of its diversity evolved in the hundreds of thousands of years before domestication. Here, by generating a database of haplotypes from the NAM founders, we try the first implementation of the PHG to impute samples within the structurally diverse maize species.

This paper asks whether the PHG, implemented as a database of haplotypes and an imputation platform, can address the issues of read mapping, haplotype library completeness, and suitability for genomic and breeding applications. The PHG pangenome representation and alignment processing should help deal with read mapping issues, which we tested with GBS and WGS data. Haplotype libraries are very useful in imputing across breeding programs (e.g., FILLIN), and here we test if the haplotype diversity from the NAM founders can be leveraged through the PHG for genotyping across the NAM RILs and a diverse population. Finally, we compare the PHG imputed genotype calls with known genotype benchmarks for each population to assess its utility in general breeding or genomic analysis.

## Results

### Representation of the pangenome in the PHG

A Practical Haplotype Graph database was constructed from 27 diverse inbred lines: the 26 parents of the Nested Association Mapping panel (McMullen et al., 2009; *NAM Genomes Project*, 2020) and B104 (reference TBA). The database consists of 71,354 reference ranges: regions from physical intervals of the B73 AGPv5 genome sequence. The edges for each reference range are defined by the starting and ending points of the gene annotations for the B73 assembly, resulting in alternating genic and intergenic reference ranges. These reference ranges allow for the identification of the haplotypes in the pangenome through the sequence aligned to them from each of the 26 other assemblies. In some cases, reference range breakpoints (i.e., edges of genes) could not be aligned from the non-B73 to the B73 assembly, likely due to presence-absence variation.

On average, non-B73 assemblies had sequences aligned to ~87% of the B73 reference ranges (Fig.1). 80% and 69% of intergenic and genic ranges were present in all taxa (Fig.2). When comparing each NAM founder’s background, it is apparent that tropical and sweet types have more missing haplotypes relative to the selected B73 reference. However, the sequence contained in the database does represent an average of 99% of each assembly (Fig.3), indicating that almost the complete sequence of them is represented in the stored haplotypes. Additionally, the genome coverage is not affected by the background. The assembly for Oh7B is a relative outlier due to a translocation of sequence between Chr9 and Chr10 (Albert et al., 2010). Our pipeline, which aligns pairs of equivalent chromosomes, misses those haplotypes. However, this translocation is not present in the Oh7B lineage used in breeding programs nor the creation of the NAM RILs. The next version of the PHG database will address this translocation to represent the NAM version of Oh7B.

**Fig.1.**
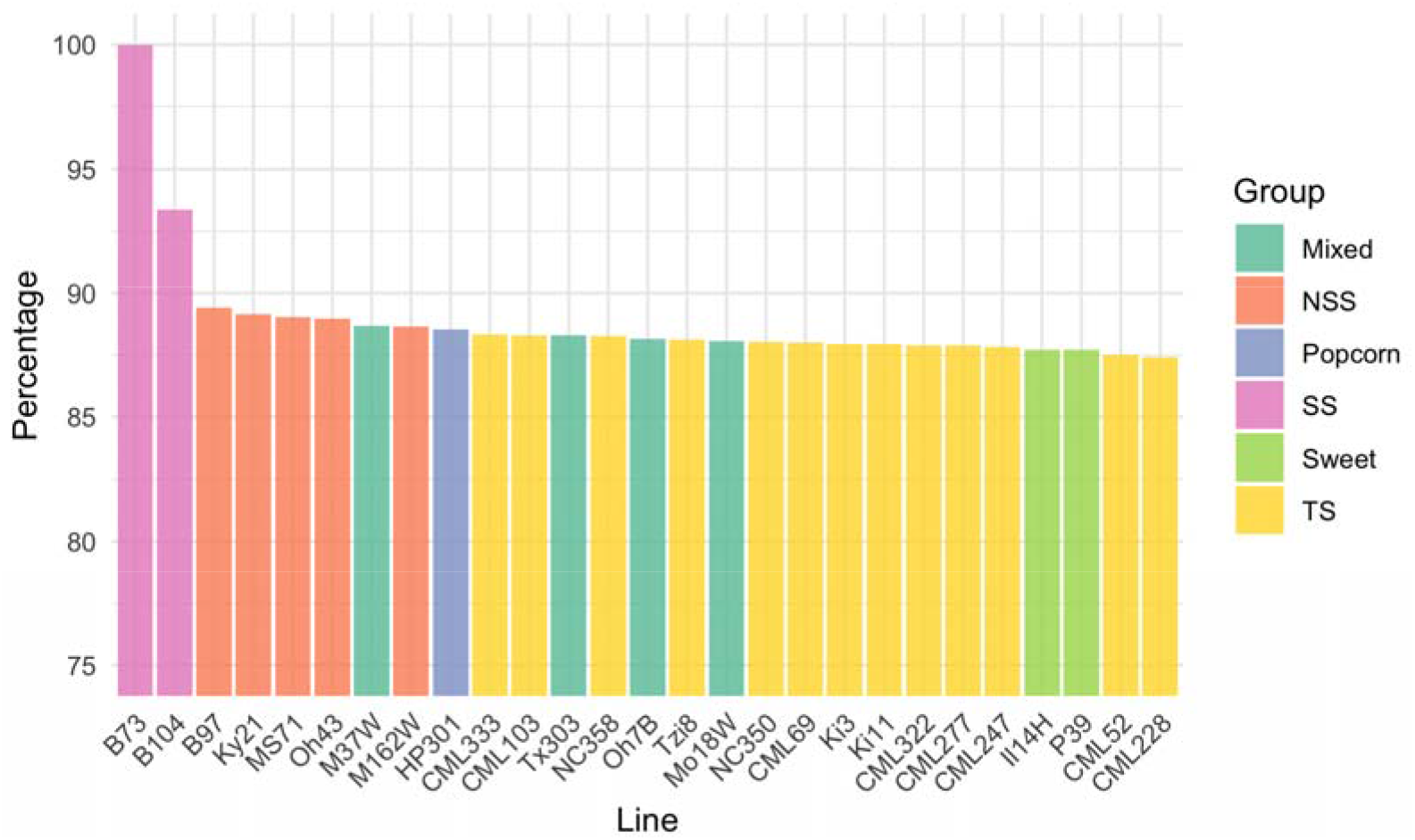
Percent of defined reference ranges with identified haplotypes in the assemblies stored in the database. B73 is at 100 as it is the assembly defining the haplotype regions.

**Fig.2.**
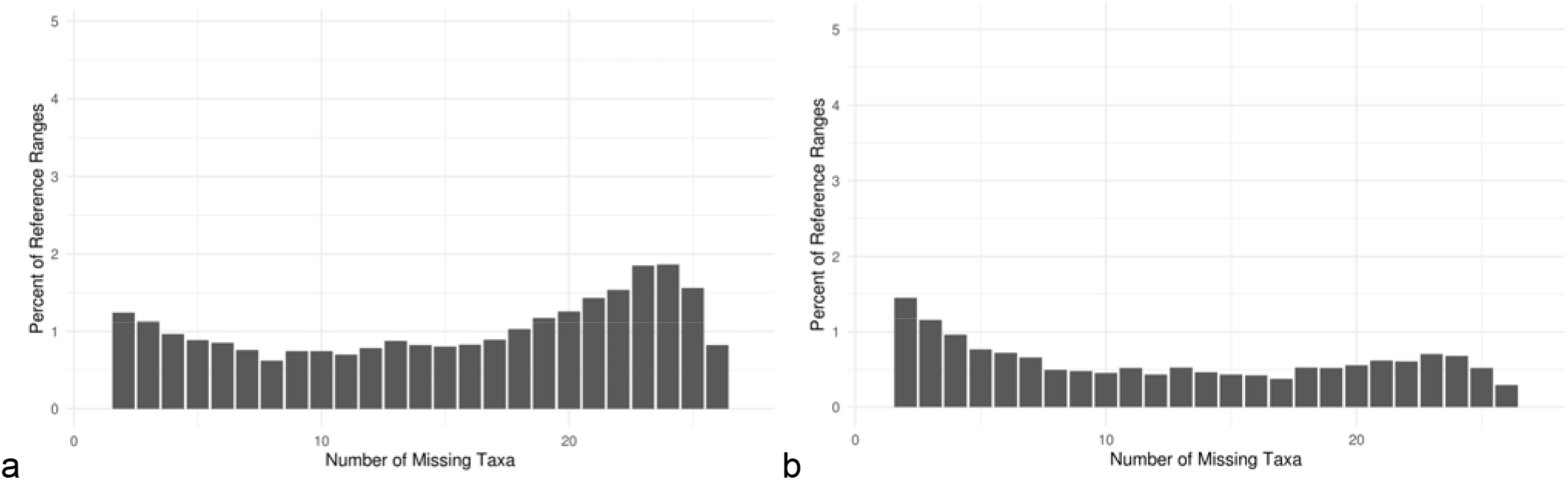
Reference range haplotype missingness. For genic (a) and intergenic (b) regions. A large portion of the reference ranges is found across all taxa, with a small portion of them being private to a subset. Bars at 0, not shown, represent 69% and 80% of genic and intergenic ranges being found across all taxa, respectively.

**Fig.3.**
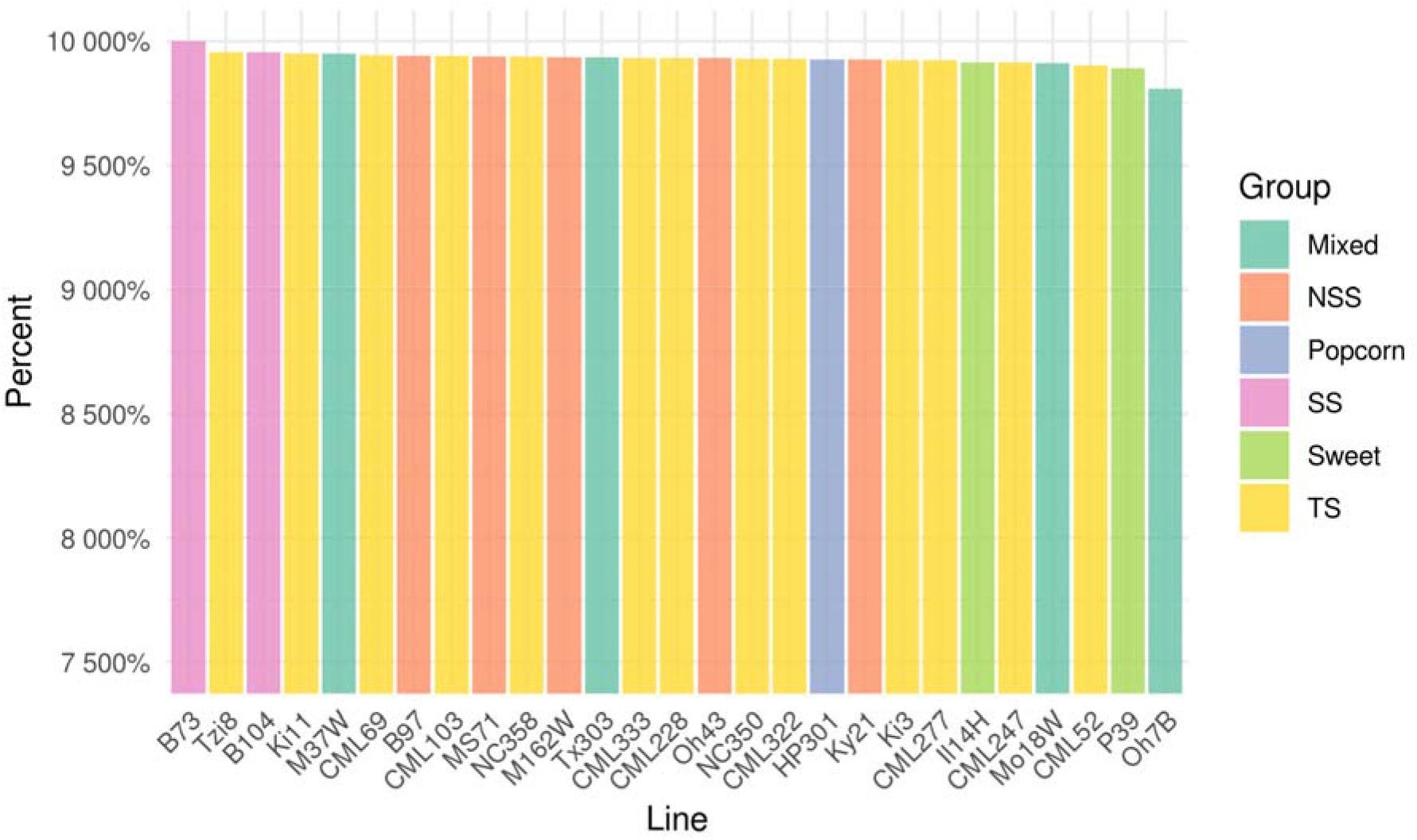
Percent of total assembly sequence contained in the PHG database for each assembly. Note that regardless of haplotypes not being identified for several reference ranges, the final length of the sequence contained is not severely impacted, as “novel” sequences potentially found surrounding the missing haplotypes are included in the adjacent reference range.

Comparing each assembly to B73, we identified a median of 1M genic SNPs and 8M intergenic SNPs, for a combined total of 43.1M SNPs over all the assemblies in the database (Fig.4). B73 genic and intergenic divergence agrees with known pedigree backgrounds.

**Fig.4.**
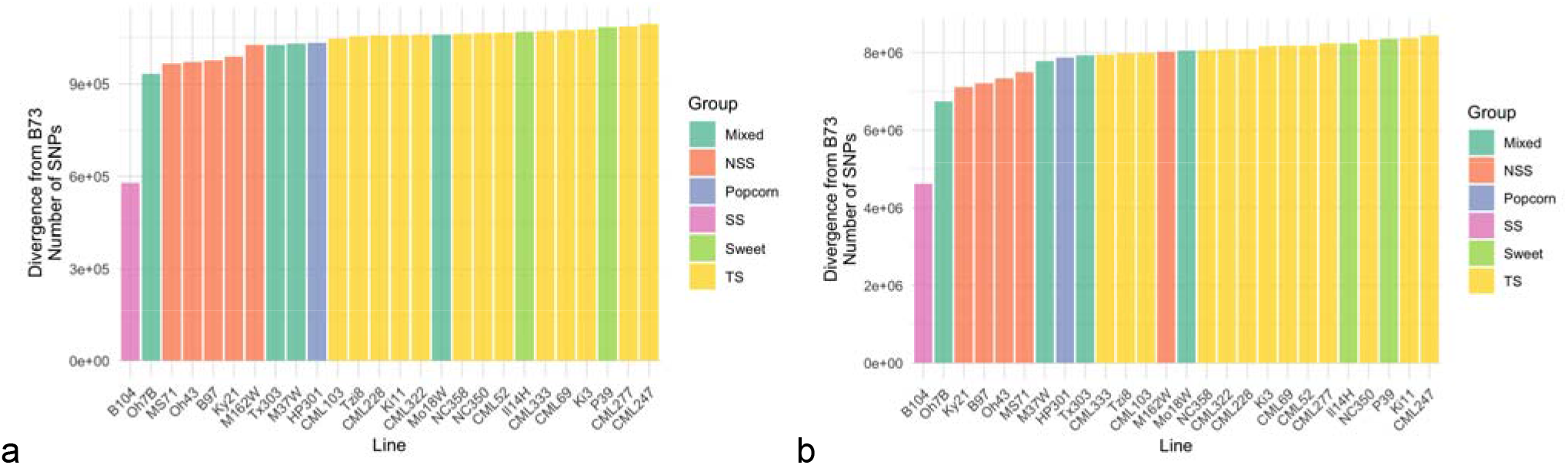
Number of SNPs identified on the haplotypes for each assembly’s reference ranges, over genic (a) or intergenic (b) regions. Known genetically close varieties to B73 show the least number of SNPs, while more distant ones have higher numbers.

### Genotyping and imputation of related lines using GBS

As an initial test for the maize PHG, we mapped GBS reads from 4705 accessions of the NAM RIL population. The GBS reads are generated with earlier Illumina/Solexa sequencers having ~70bp in length with extremely low coverage (more modern technologies would produce longer and more reads). These were mapped to the pangenome to impute haplotypes and generate SNP calls. To evaluate the imputation accuracy against existing results, we compared the imputed SNPs to 1,106 legacy SNPs (McMullen et al., 2009) and observed an average error rate of 0.8% (Fig.5a and 5b). The families with the larger error rates have known residual heterozygosity, CML52 at 1.4%, Tzi8 at 1.7%, and CML228 at 1.8%. The error rate by position appears evenly distributed throughout the chromosomes (Fig.Sup.1), except for three positions in chromosomes 4, 7, and 8, where all families have higher than average error rates for a subset of the taxa.

**Fig.5a.**
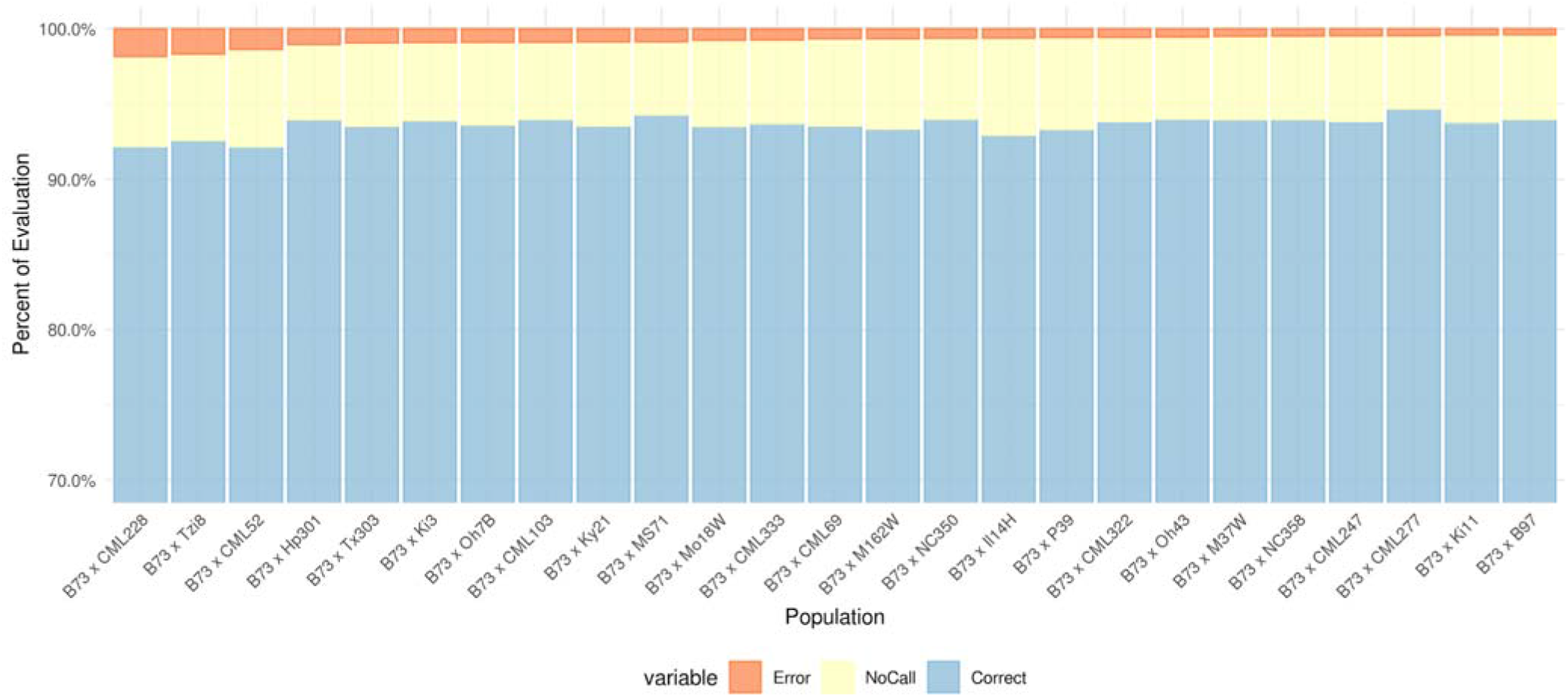
Evaluation of imputed sites for each NAM RIL over 1107 uplifted legacy SNPs used as a benchmark. Evaluations are grouped by family for simplified viewing. Bars are sorted by percentage of errors. No calls are the product of residual heterozygosity, low density and missing data in benchmark SNPs, or small unresolved breakpoints between the GBS reads.

**Fig.5b.**
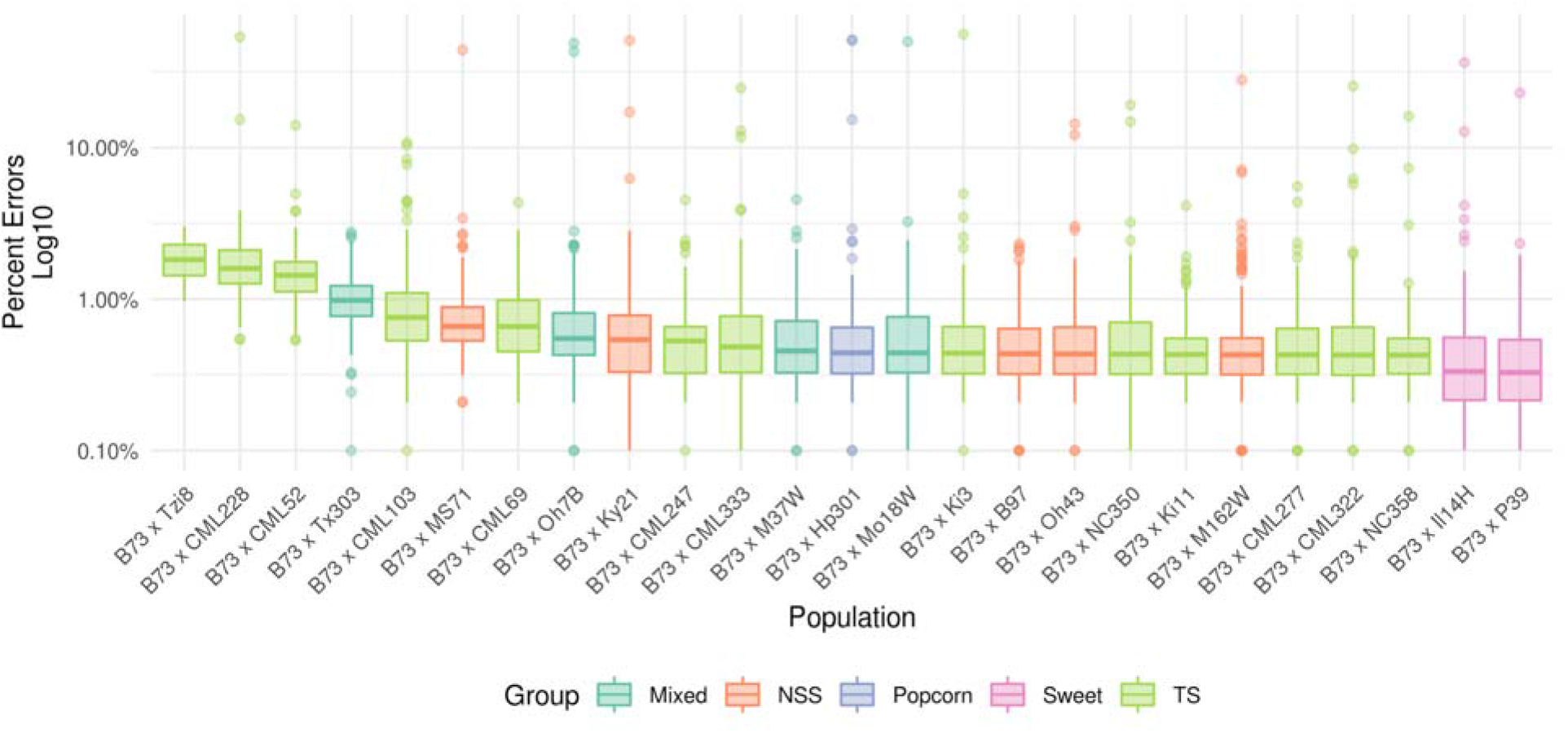
Per family accession error. Colored by family background. Y-axis is log10 scaled. The presence of a few outliers in individual families does not increase the average family error rate. The background does not affect imputation accuracy.

### Imputation of unrelated lines using GBS and WGS

To assess the PHG’s ability to impute samples indirectly related to the included haplotypes, we tested it on the Goodman Association Panel population, which includes diverse global breeding lines (Flint-Garcia et al., 2005). After mapping both GBS reads and WGS reads for a subset of them, we imputed haplotype paths and called SNPs as with the NAM RILs. The uplifted 600K Axiom SNP array was used as a benchmark; however, this benchmark is biased toward temperate and European diversity (Unterseer et al., 2014).

We first compared the SNPs imputed by the PHG from GBS reads. We were able to identify an average error rate of 3.4% (Fig.6). For taxa represented in the PHG, we saw an average of 0.7% error rate, while for taxa not found in the database, the average error rate was 8.3%. F7 and EP1 had the highest error, at 10% and 12%, respectively. This was not unexpected, as they represent taxa with a European background that is not well represented in the current pangenome (NAM parents plus B104). We saw negligible differences in the accuracies between genic and intergenic regions, having overall average error rates of 3.4% and 3.5%, respectively (data not shown).

**Fig.6.**
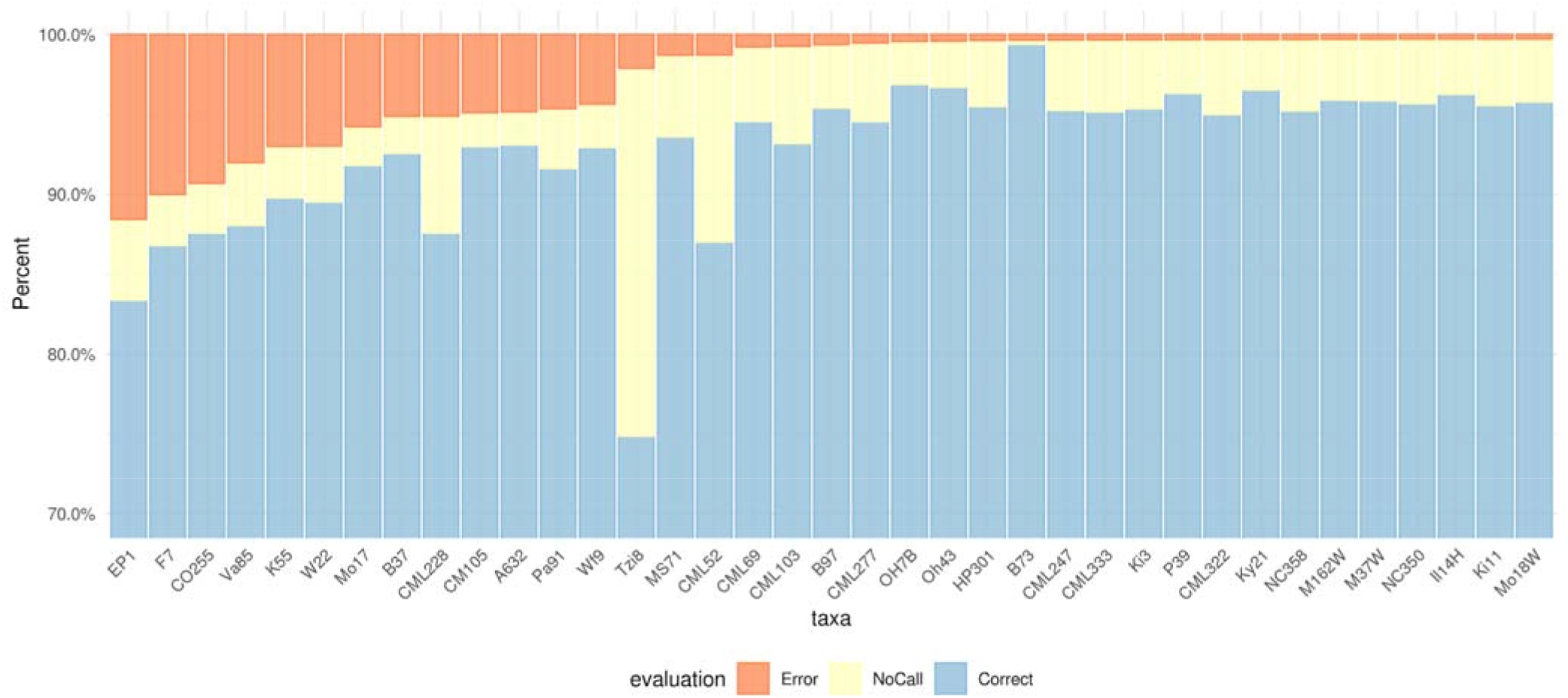
Evaluation of imputation of the Goodman Association Panel population using GBS data. Evaluated against the uplifted 600k Axiom SNP array. Blue and orange show the proportion of correct and erroneous calls, respectively. In yellow are the proportion of SNPs where the benchmark SNP was heterozygous and masked, or where the PHG does not impute that site. Bars are sorted by decreasing error-rate.

To assess the effect of having higher sequencing coverage and depth, we evaluated SNPs imputed by the PHG from WGS reads. We observed a decrease in the proportion of errors to an average of 2.2% (Fig.7), with average error rates of 0.7% and 5.3% for taxa represented or missing from the database, respectively. The average genic and intergenic error rates were 1.9% and 2.5%, respectively. The largest effect, albeit still small compared to when using GBS reads, was on the number of imputed sites; the average percentage of missing sites decreased from 4.5% to 2.2%.

**Fig.7.**
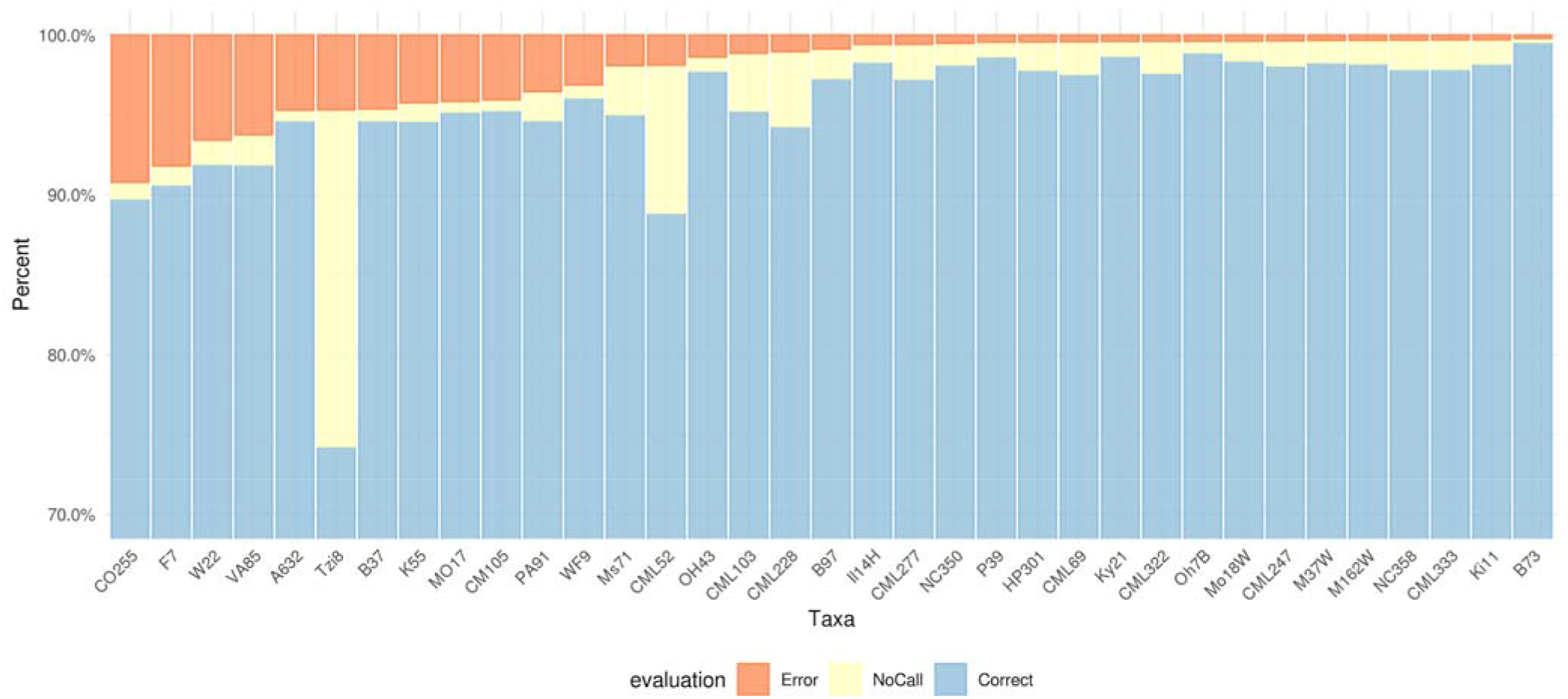
Evaluation of imputation for a subset of the Goodman Association Panel population through WGS data. Evaluated against the uplifted 600k Axiom SNP array. Blue and orange show the proportion of correct and erroneous calls, respectively. In yellow are the proportion of SNPs where the benchmark SNP was heterozygous and masked, or where the PHG had not imputed that site. Bars are sorted by decreasing error-rate.

### Assessing causes that influence error rates

To identify the causes driving the errors, we analyzed four potential causes: missing haplotypes, recombination rate, minor allele frequency, and haplotype read counts.

#### Missing haplotypes

We compared the clustering of the errors across the genome to identify whether errors are due to missing haplotypes or rare alleles (Fig.8). When compared with a set of SNPs with randomized positions, we observed that errors are clustered in longer runs than expected at random. This effect is stronger when imputing taxa not represented in the database. This indicates that the current pangenome is missing haplotypes over a small set of reference ranges, which produce most of the errors. The inclusion of rare alleles not present in the database (~44,000 sites) has no effect (data not shown). The main source of error in the current pipeline is from the absence of important haplotypes, not rare alleles.

**Fig.8.**
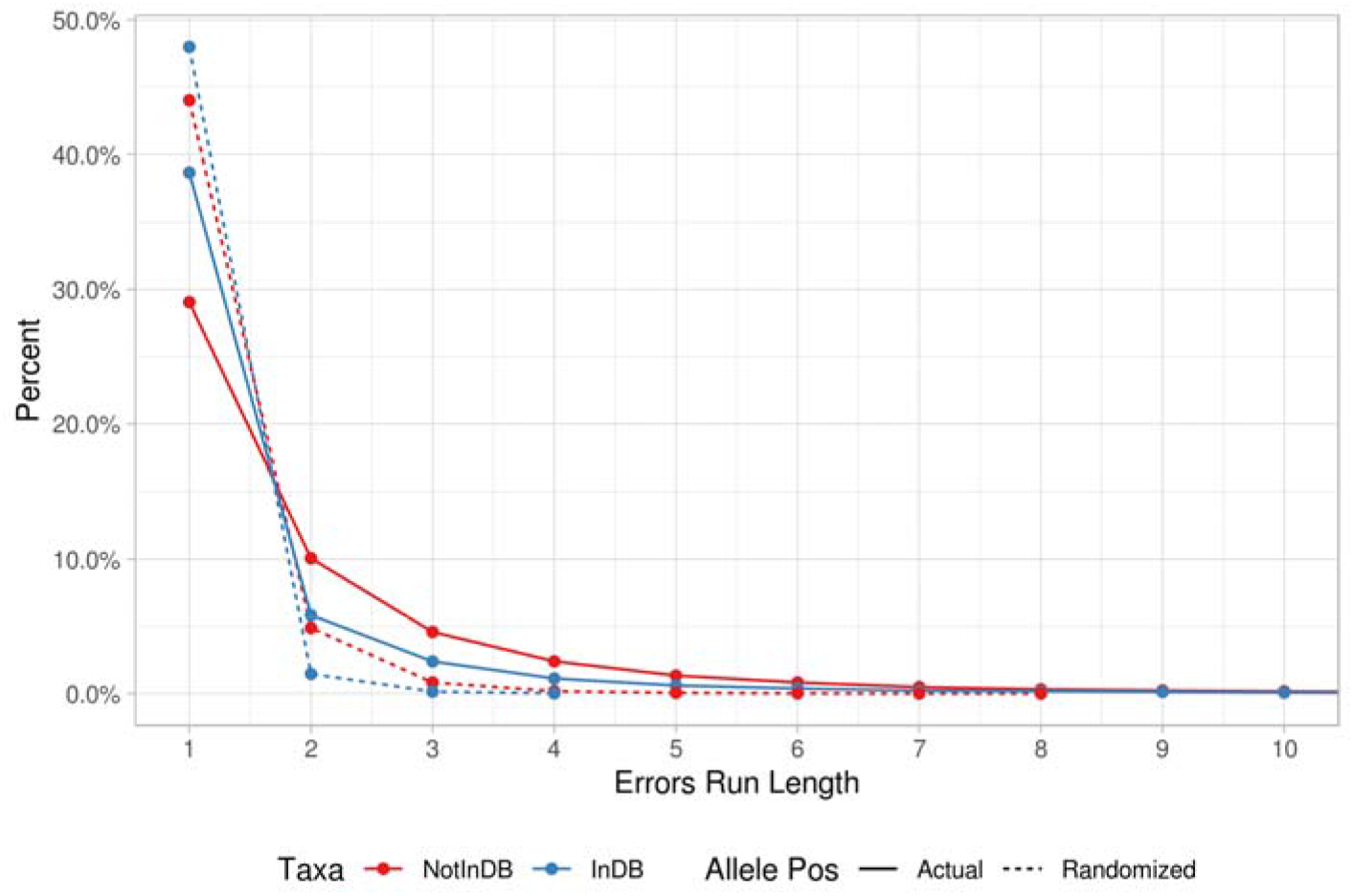
Runs of erroneously imputed sites. Imputing taxa present in the PHG haplotype database permits for longer runs of correctly imputed SNPs, with taxa not present in the PHG database having longer stretches of incorrectly imputed SNPs. Randomized allele positions are sampled randomly from 30% of the sites for each taxon and chromosome. Taxa were grouped by whether they were represented in the pangenome database (blue) or not (red).

#### Recombination rate

Another potential source of errors arises when samples have recombination within a reference range, resulting in two or more haplotypes. This is likely due to different rates depending on location in the genome, given that that recombination in maize occurs in a frequency that can vary within nearly two orders of magnitude along chromosomes (Rodgers-Melnick et al., 2015). Higher recombination rates should decrease the PHG’s ability to represent haplotypes accurately. However, higher recombination rates occur near genic areas. Because gene boundaries were used as haplotype breakpoints, the effect of recombination rate on errors could be minimal. We tested if the recombination rate was correlated with the error rate in 100 equally sized bins of the recombination rate (Fig.9). Error rates for taxa not represented in the PHG database appear correlated with an increased recombination rate. As expected, taxa within the PHG database do not show the same pattern. The inclusion of rare alleles in the analysis did not change the effect. While minor allele frequency (MAF) correlates with error rate (Fig.Sup.2a), MAF is only weakly associated with recombination rates (Fig.Sup.2b). This shows that the errors are also partially driven by novel haplotypes derived from recombination. Increasing the haplotype sampling or decreasing the reference range lengths in high recombination areas could alleviate this problem.

**Fig.9.**
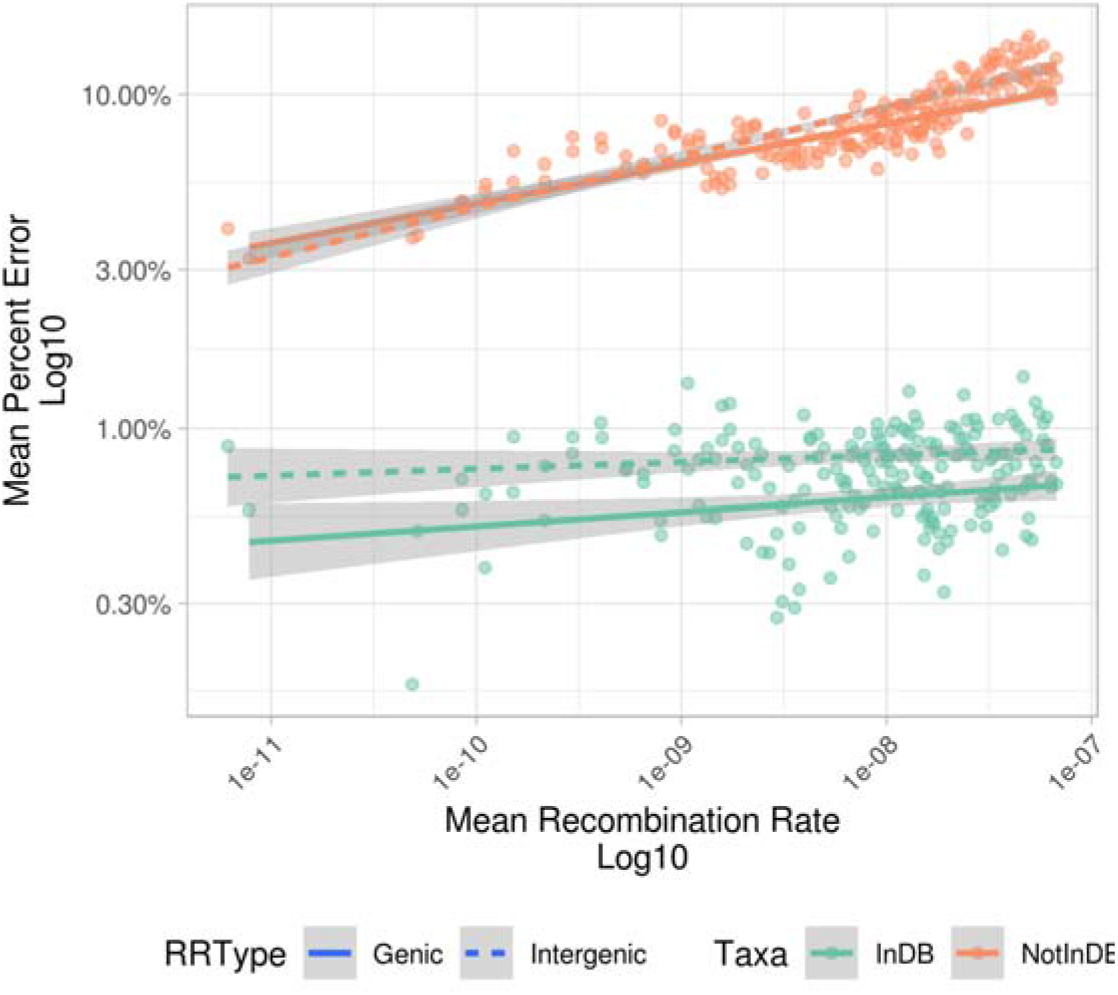
Binned analysis of per-site error rate by recombination rate. Averaged over 100 bins of equal size after the data was sorted by recombination rate. Taxa were grouped by whether they were represented in the pangenome database (green) or not (orange).

#### Haplotype read counts

A large number of duplicated regions in the maize genome complicates accurate read mapping. The PHG pipeline addresses this by keeping read mappings only when they map within a single reference range. The identification of a specific haplotype within a reference range should increase with haplotype coverage. To assess the sensitivity of the PHG to read mapping coverage, we tested the effect of the number of reads mapped to the error rates on the imputed haplotypes over 100 bins of increasing mean number of mappings (Fig.10). Error rates for taxa not in the PHG database decrease slightly with an increasing mean number of reads. As expected, this effect is smaller for taxa in the database. In both cases, the effect was higher in genic ranges.

**Fig.10.**
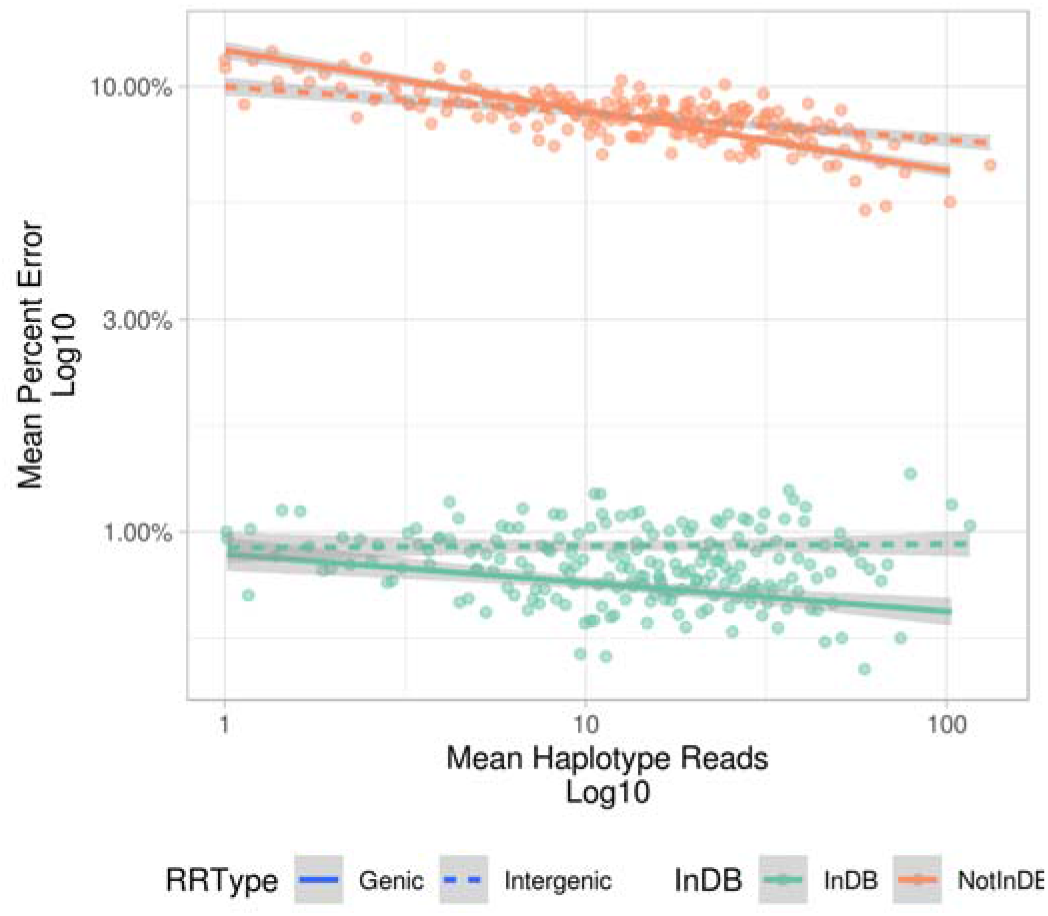
Per-haplotype error rate by mean haplotype read count. Binned haplotype error rates were calculated over 100 equally-sized bins of an increasing number of reads mapped. The analysis is shown separately by genic or intergenic reference range type and taxa grouped by whether they were represented in the pangenome database (green) or not (orange).

With the sequence context for over 40 million SNPs, this maize PHG database occupies a modest 83GB of disk space, being effectively only slightly larger than the genomes it represents. For comparison, a VCF file for the same taxa number, without haplotype data or mapping information, requires over 1TB of space. By leveraging the diverse NAM founders and their high-quality assemblies through the PHG, we have shown its utility in imputing related and unrelated samples. While the current database cannot perform as accurately on accessions with unrepresented haplotypes, we expect the future addition of a relatively small set of assemblies will address this issue. We expect the maize PHG will enable the community to more efficiently and effectively identify the genetic mechanisms underlying the control of many more phenotypes.

**Table 1:** Populations genotyped by the PHG

| Population | Inbreeding | Diversity | Sequencing Depth |
| --- | --- | --- | --- |
| NAM RILs | High | Low | Low |
| Goodman Association Panel GBS | High | Moderate | Low |
| Goodman Association Panel WGS | High | Moderate | High |

**Table 2:** Summary Main Genotyping results

| Population Panel |  | NAM - 1k legacy SNPs | Goodman Association Panel - 600k Axiom SNP chip |  |
| --- | --- | --- | --- | --- |
| Taxa in PHG database\Data type |  | GBS | GBS | WGS |
| Average PHG benchmark agreement* | In DB | 99.2% | 99.3% | 99.3% |
|  | Not in DB |  | 91.7% | 94.3% |

\*Considering only imputed and not heterozygous sites in the benchmark.

## Discussion

Here we have presented a first maize PHG, a pangenome haplotype database, and a genotyping pipeline that allows the accurate imputation and reconstruction of whole-genome haplotypes and genotyping with minimal information. By parting from the concept of a closed pangenome, where a limited number of taxa can represent virtually all of the haplotypes found in a species, we leveraged the NAM assemblies in the PHG to create a novel genotyping and imputation tool. The maize PHG will thus facilitate community genotyping efforts by achieving higher accuracy of imputation and implementing a standardized approach that will return consistent results from low coverage sequencing.

Processing the assemblies through the PHG pipeline allows us to identify homologous haplotypes among the included assemblies for most defined reference ranges. Previous analyses of variation in intergenic (Anderson et al., 2019) and genic regions (Sun et al., 2018) show a relatively similar pattern to the results observed in this maize PHG database, having regions essentially shared at ~80% rate, with a relatively small subset of them being mostly missing in the majority of taxa. This is only a first approach, as potentially a more complete gene model set, resulting in smaller reference ranges, might better reflect the presence/absence variation of more refined haplotypes. Additionally, choosing a distinct base reference that more closely resembles the larger pangenome might also allow for the more direct dissection of otherwise less common haplotypes.

By imputing haplotypes known to be within our database, as shown with the NAM RILs, we demonstrate the pangenome’s ability to act as a useful target for read mapping. Moreover, the pipeline functions as an effective tool to process these mappings and accurately identify the underlying haplotypes from which those reads are derived. This is of particular interest for the maize and general plant communities, due to the frequently high repetitive nature of their genomes, where the unambiguous mapping of short sequences continues to be a challenge for accurate and large scale genotyping.

Assessing the genotyping accuracy on the diverse Goodman Association Panel population, we also show that this current maize PHG database, while effective, is missing a relatively small but significant portion of the broader maize pangenome. As opposed to rare alleles, this lack of haplotypes was the main source of errors, as evidenced by the clustering of errors in a subset of the haplotypes and over those regions with a high recombination rate. This is particularly the case for samples of European backgrounds, which are not well represented in the implemented database. We expect an increase in haplotype diversity within the PHG database, by including additional diverse genomes such as those from CIMMYT maize lines, will allow the PHG to be a one-stop solution for the accurate genotyping of most maize lines.

This first maize PHG database consists of a pangenome haplotype database from 27 high-quality maize assemblies. It includes the information on the mapping and imputation of over 5,000 NAM RILs and Goodman Association Panel accessions. We expect that this maize database, along with the broader PHG pipeline, through its ability to generate accurate genotype calls using either GBS or WGS reads, small computational footprint, and ease of transferability will help the maize community achieve accurate and inexpensive genotyping, enabling more analyses to discover the genetic basis of many more phenotypic traits.

## Materials and Methods

### Building the PHG

We populated a maize PHG database (Bradbury, 2020) to store the pangenome’s information and keep track of reference ranges, haplotypes, mappings, and paths imputed. Briefly, the PHG is a relational database that divides the reference genome in ranges, subdivides other pangenomes in similar ranges, and allows for rapid and efficient storage and retrieval of information about the reference ranges, component haplotype IDs, and paths. A Java and R API have been developed for the PHG software package to implement and interact with this pangenome database (Bradbury, 2020; Bradbury et al., 2007).

To populate the maize PHG database, we generated a pangenome by leveraging the high-quality assemblies for the 26 diverse NAM parents (*NAM Genomes Project*, 2020) and B104 (reference TBA). To define the pangenome haplotypes, we took a three-stage approach. First, we selected the B73 RefGen_v5 (*MaizeGDB*, 2019) as the base reference. This allows for a direct comparison with existing genotyping platforms and results. Second, we defined the haplotype regions by choosing a set of reference ranges. We selected the B73 RefGen_v5 gene regions (Zm00001e.1) to allow for haplotype boundaries that are: clearly defined, of biological significance, and more likely to be conserved. We used the 35,677 genic regions as breakpoints for the definition of the 71,354 B73 haplotypes. Third, we used Mummer4 (Marçais et al., 2018) to identify the haplotypes in those regions on each assembly. Briefly, the assemblies were divided into individual chromosomes. Each chromosome was aligned against the B73 RefGen_v5 equivalent using the nucmer program of Mummer4. Testing values for the -c parameter between 150-500, we set the value to 250, to find a balance between speed and the length of maize exons (Haberer et al., 2005; *MaizeGDB*, 2020a). These alignments were then processed, with each reference range in each assembly having a haplotype ID assigned and stored in the database. Insertions or deletions found within each reference range are stored as part of the identified equivalent haplotype, regardless of their size. Reference ranges for taxa that do not produce an alignment are left empty (Bradbury, 2020).

### Alignment to Pangenome

To evaluate the ability of the PHG to impute haplotypes, we utilized the GBS reads originally generated for the NAM (Rodgers-Melnick et al., 2015) and the Goodman Association Panel (Romay et al., 2013). Additionally, WGS paired-end reads (Bukowski et al., 2018) were obtained for a subset of the Goodman Association Panel samples (SRA study accession number SRP108889), which had also been genotyped with the Axiom 600k SNP array (Unterseer et al., 2014). This allows us to compare the ability of the PHG to impute haplotypes to a distinct and diverse set of sequencing data. CO125 was removed from the analysis as its genotyping data suggests a sample mixup in its sequencing.

The reads were mapped to an index of the pangenome generated by minimap2 (Li, 2018) (Ver. 2.17-r941). We set parameters -k 21 -w 11 -I 90G to reflect the recommended short read alignment kmer size and the necessity of having the complete pangenome sequence loaded into memory at once. Failing to set -I large enough to fit the whole pangenome in RAM returns a multi-part index, which produces poor mapping processing results. The read mappings made use of the short read heuristics (-k21 -w11 --sr --frag=yes -A2 -B8 -O12,32 -E2,1 -r50 -p.5 -N25 -f1000,5000 -n2 -m20 -s40 -g200 -2K50m --heap-sort=yes). To maximize the likelihood of getting read mappings to all matching haplotypes, we modified the flag -N to produce 25 secondary mappings. The mappings were then processed through and stored in the PHG database. Briefly, only edit distance optimal alignments are considered, reads that map to multiple reference ranges are discarded, and reads that map to specific haplotypes are identified among the read mappings for the haplotypes within the reference range.

### Imputation evaluation

Once the read mappings are loaded into the PHG database, imputation is done by finding a path through the haplotype graph using the BestHaplotypePathPlugin in the Java PHG API. Briefly, the read mappings are used to count the number of times reads mapped to each haplotype. Using these counts, and the transition probabilities between adjacent haplotypes, an HMM algorithm finds the most likely haplotype path through the graph. The parameter minReads is set to 0 so that the algorithm imputes haplotypes for all reference ranges, including those that have no reads mapping to them. The resulting imputed paths are stored in the PHG database. Finally, all SNPs within imputed haplotypes are exported as a VCF file (PathsToVCFPlugin). This generates a 1TB VCF with >42M sites for the 4,705 NAM accessions. A similar process was followed to impute and generate the SNP calls for the Goodman Association Panel’s taxa.

An SNP benchmark set was defined for the NAM RILs and Goodman Association Panel population samples to evaluate imputation accuracy. For the NAM population, the 1,144 legacy SNP set (McMullen et al., 2009), NAM_map_and_genos-120731.zip, was obtained (*Panzea*, 2009) and uplifted from AGPv2 to v5 in a two-step approach. First, from the original v2 to v4, we used CrossMap (Zhao et al., 2014) and the corresponding chain file (*ENSEMBL*, 2020). These were then uplifted from v4 to v5 using the liftover_vcf pipeline (https://github.com/qisun2/liftover_vcf) and a v4 to v5 chain file (*MaizeGDB*, 2020b). This returned 1,106 variant sites uplifted to v5 coordinates, which we then used to compare our imputation results. For the Goodman Association Panel, the 600 K (Unterseer et al., 2014) Axiom SNP array was uplifted from AGPv4 to v5 coordinate equivalents through Crossmap and utilized as a benchmark.

Custom code was written to evaluate the accuracy of the imputed SNP calls by comparing them to the benchmarks. In short, the imputed VCF files were intersected with the Axiom SNP sites using bedtools (Quinlan & Hall, 2010). These VCF files were then loaded into R (R Core Team, 2018) through the SNPRelate package (Zheng et al., 2012). Heterozygous genotypes in the benchmark were set as missing, as the haplotypes are expected to be homozygous, and the current pipeline imputes homozygous SNPs. Then, the two sets of SNPs were matched to the equivalent taxa, and the correspondence of reference or alternate allele calls were evaluated as correct or incorrect if they agreed or not. Error rates are calculated as the number of incorrect allele calls divided by the total number of correct *and* incorrect calls. For sites on taxa where the PHG makes no allele call, or where the benchmark has a heterozygous allele, the evaluation is set as NoCall. To identify the proportion of errors between genic and intergenic regions, the GenomicRanges package (Lawrence et al., 2013) was used to identify the SNPs as found within either set of regions, and the data.table (Dowle & Srinivasan, 2019) package was then used to summarize each region type’s calls.

### Assessing causes that influence the error rate

#### Evaluating runs of errors

SNP evaluations sorted by position were processed through the rle function (R Core Team, 2018) of R to get the run-length of error calls on each of the chromosomes for each taxon. These runs of errors were then analyzed by whether the taxa were represented in the pangenome database or not. A random set of 30% of the SNP evaluations for each taxon and chromosome were selected, effectively randomizing the SNP positions, on which we then calculated the run-length of errors. The frequency of each of the run-length of errors was then calculated for each category.

#### Comparison of error rate vs. recombination rate

Recombination rates on the NAM population were obtained from (Ramstein et al., 2020). These were uplifted from v4 to v5 through Crossmap. The uplifted values were then matched to the evaluated SNP calls. This combined data set was then sorted by the recombination rate. The mean recombination and error rate were calculated over 100 bins of equal size number of SNPs.

#### Comparison of error rate vs. minor allele frequency

Minor allele frequencies were obtained from the Axiom 600k SNP array from the taxa under analysis through the snpgdsSNPRateFreq of the SNPRelate package. These frequencies were matched to the evaluated SNP calls. One hundred bins of equal length were then created after sorting by increasing recombination rate or minor allele frequency. The means for error rate, recombination rate, and MAF were then calculated on each bin.

#### Comparison of error rate vs. read counts

Imputed haplotypes for each sample were obtained from the PHG database through the pathsForMethod function of the rPHG package (Monier et al., 2019). For each taxon, read mappings were obtained from the PHG database through the readMappingsForLineName, also of the rPHG package. The mappings were then matched to the haplotypes imputed for each taxon. The mean error rate and mean read count were calculated for the haplotypes over each reference range by the sample representation status in the pangenome database and by reference range type. The calculated values over these four categories were then sorted by the mean number of reads and analyzed over 100 bins of equal size. The mean number of reads and the mean error rate were calculated for each bin.

The maize PHG database is publicly available through the Buckler Lab webpage: https://www.maizegenetics.net/post/the-first-maize-phg-database-now-available The code for these analyzes is mostly written in R and made available at: https://bitbucket.org/bucklerlab/p_maizephg

## Acknowledgments

This material is based upon work supported by the USDA-ARS, NSF Research-PGR Grant No. IOS-1822330, NSF Postdoctoral Research Fellowship in Biology under Grant No. IOS-1906619, Bill and Melinda Gates Foundation, and a CONACYT-I2T2 scholarship for graduate studies.

---

**Fig.Sup.1.**
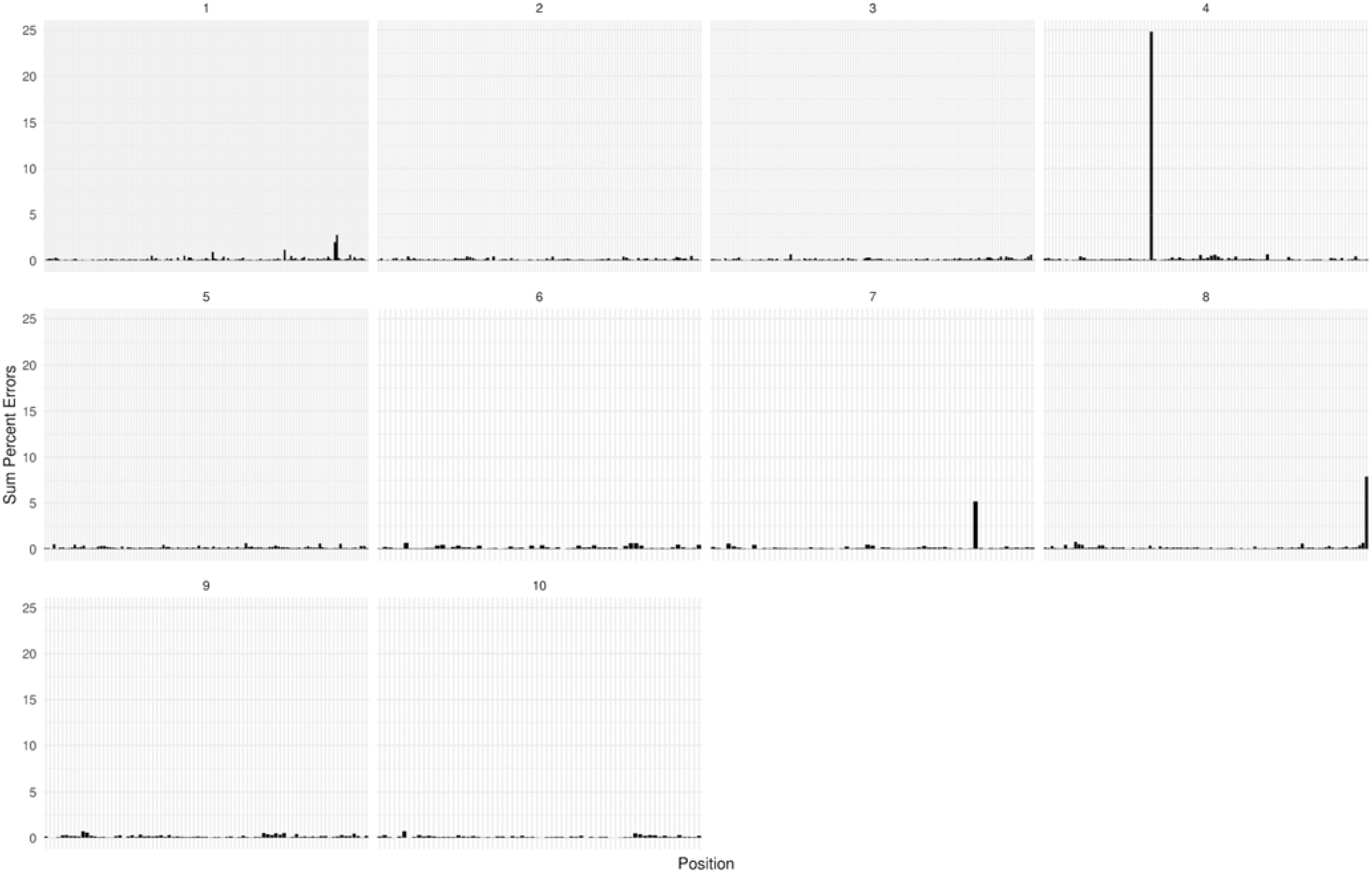
Sum of percent errors by chromosomal position. Some positions stack up a large proportion of the individually small errors across families (e.g., chromosomes 4, 7, and 8). Not shown here, some positions concentrate a majority of the errors of individual families (e.g., chromosome 6 for CML228, chromosome 10 for CML52)

**Fig.Sup.2.**
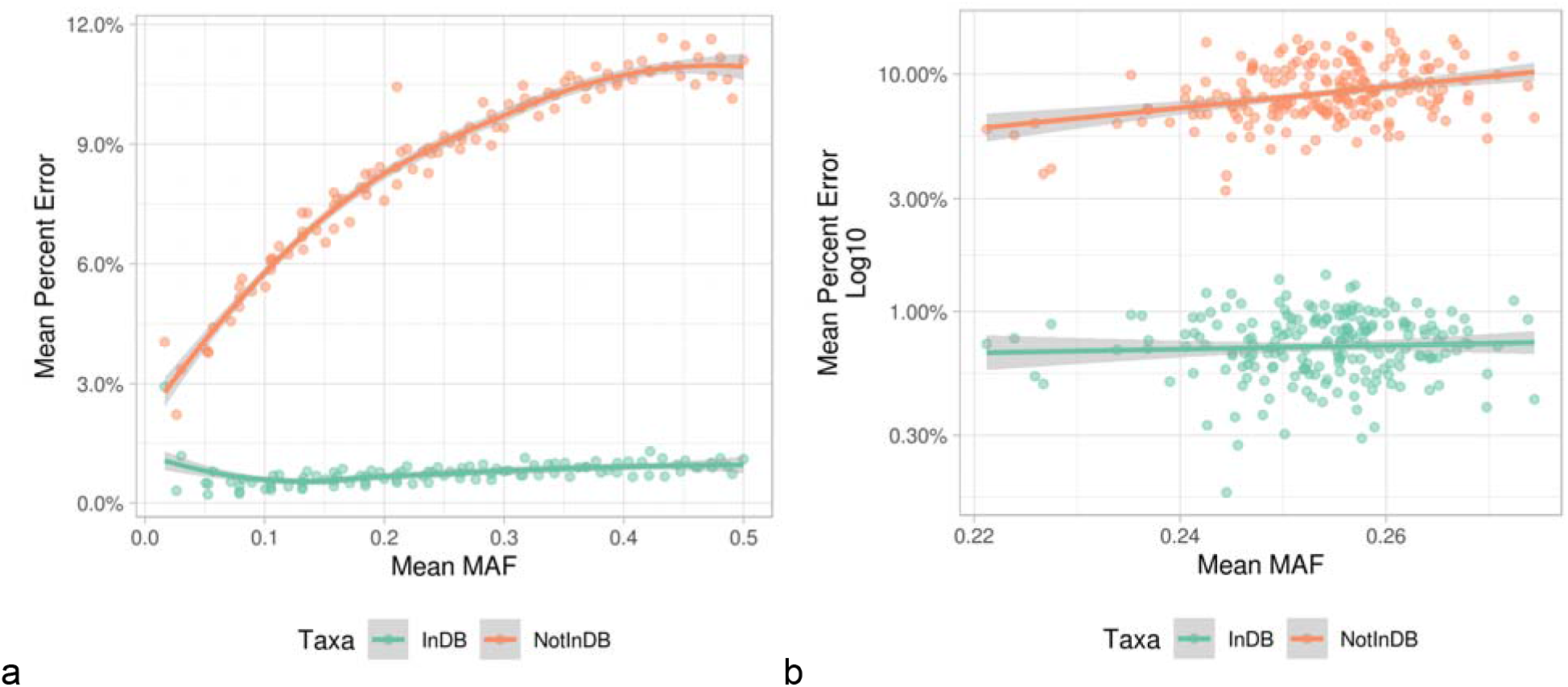
Binned analysis of the per-site error rate. Effect of MAF over error rates. Averaged over 100 bins of equal size after the data was sorted by minor allele frequency (a) or recombination rate (b). Taxa were grouped by whether they were represented in the pangenome database (green) or not (orange).

## Notes

https://www.maizegenetics.net/post/the-first-maize-phg-database-now-available

### Competing Interest Statement

The authors have declared no competing interest.

### Summary of Updates

Dating of second genome duplication. VG reference corrected.

